# Phenotypic functional activities of monocyte change during crosstalk with breast cancer cell and enhancing effect of metformin of IFN-γ-associated antitumor cytokine production

**DOI:** 10.1101/2020.02.14.949412

**Authors:** Zoheir Dahmani, Lynda Addou-Klouche, Florence Gizard, Sara Dahou, Aida Messaoud, Nihel Chahinez Djebri, Mahmoud Idris Benaissti, Meriem Mostefaoui, Hadjer Terbeche, Wafa Nouari, Marwa Miliani, Gérard Lefranc, Anne Fernandez, Ned J. Lamb, Mourad Aribi

## Abstract

**Background:** Immune activities of monocytes (MOs) can be altered within the microenvironment of solid malignancies, including breast cancer. Metformin (1,1-dimethylbiguanide hydrochloride, MET), has been shown to decrease tumor cell proliferation, but its effects have yet to be explored with respect to the crosstalk between monocytes and breast cancer cells. Here, we investigated the effects of MET on overall phenotypic functional activities of autologous MOs during the interplay with primary breast cancer cells.

**Methods:** Human primary breast cancer cells were either cultured alone or co-cultured with autologous MOs before treatment with MET.

**Results:** MET downregulated both breast cancer cell proliferation and the ratio of phosphorylated Akt (p-Akt)-to-Akt in breast cancer cells. Additionally, we observed that, in the absence of MET treatment, the levels of LDH-based cytotoxicity, catalase, intracellular free calcium ions (_if_Ca^2+^), IL-10 and arginase activity were significantly reduced in co-cultures compared to those of MOs cultivated alone whereas levels of iNOS were significantly increased (for all comparisons, *p* < 0.05). In contrast, MET upregulated breast cancer cell LDH-based cytotoxicity levels when co-cultured with MO. MET also induced upregulation of both the inducible enzymatic activity of nitric oxide synthase (iNOS) and arginase activity in MO cells and co-culture systems, although these differences did not reach significant levels for iNOS activity (*p* > 0.05). MET greatly decreased phagocytic activity in isolated MOs while inducing a robust increase of catalase activity in co-culture systems and of superoxide dismutase (SOD) activity in MOs, but not in MOs co-cultured with breast cancer cells. MET strongly upregulated the levels of _if_Ca^2+^ in co-culture systems and IFN-γ production in both isolated MOs and co-culture systems. Moreover, MET treatment markedly downregulated IL-10 production in MOs, while inducing a slight increase in co-cultures (*p* > 0.05).

**Conclusions:** Our results show that the phenotypic functional activities of MOs change when co-cultured with primary human breast cancer cells. Furthermore, treatment with MET induced enhancing effects on the production of antitumor cytokine IFN-γ and _if_Ca^2+^, as well as cytotoxicity during breast cancer cell-MO crosstalk.

**Novel Highlights include:**

- First analysis of the anti-tumoral effects of Metformin on primary human breast cancer cells and the crosstalk with autologous monocytes.
- Phenotypic functional activities of monocytes change during their interplay with breast cancer cells, which is improved by upregulation of IFN-γ after Metformin treatment.
- Metformin induces downregulation of phosphorylated-Akt1/2-to-Akt1/2 ratio in breast cancer cells.
- Metformin downregulates phagocytic capacity of monocyte from breast cancer patients.

## Introduction

Breast cancer is the most commonly diagnosed cancer and a leading cause of mortality worldwide (1). Compared to other types of cancer that are considered as more responsive to immunotherapy, breast cancer has not been traditionally considered as an immunogenic malignancy (2). However, recent research has shown the relationship between immune intra-tumoral responses and breast cancer development (3). Additionally, studies reported that infiltration of immune cells within the tumor microenvironment and the presence of immunity-related gene signatures contribute to breast cancer prognosis (4,5).

The microenvironment surrounding breast cancer cells plays an important role in modulating cancer growth and progression (3). It contains several types of inflammatory cells including MOs and macrophages. MO cells represent a heterogeneous population derived from myeloid lineages (6) that are recruited from the bloodstream to the tumor site through the paracrine action of cytokines and chemokines released by breast cancer cells (7). Previous reports suggested that infiltration of MOs into the breast tumor microenvironments, in response to paracrine stimulation, correlates with poor prognosis and promotion of tumor growth, invasion and metastasis (8,9).

In light of their functional phenotypic plasticity, MOs can be targeted by several therapeutic molecules that switch them towards proinflammatory/anti-tumoral killer cells (10,11).

It is well known that insulin is an important growth factor, which plays a critical role in regulation of cell proliferation. As such, enhancing insulin sensitivity can lead to tumor growth inhibition and cell cycle arrest. Indeed, metformin (1,1-dimethylbiguanide hydrochloride, MET), an antidiabetic drug prescribed for patients with type 2 diabetes (12,13), has been reported to have a marked effect on insulin sensitivity through inhibition of the signaling pathway implicating phosphoinositol-3-kinase (PI3K) and Akt (also referred to as protein kinase B, PKB) consequently leading to decreased tumor cell proliferation (14,15). The effects of MET on breast cancer cells has also been associated with the inhibition of pro-tumoral M2-like macrophage polarization (16). In this context, we investigated for the first time the effects of MET on the overall phenotypic functional activities of autologous MOs during their crosstalk with breast cancer cells.

## Materials and Methods

### Materials

Unless specified, all materials including (MET), were obtained from Sigma-Aldrich (Sigma Chemical Co., St. Louis, USA).

#### 1. Study design

Tumor epithelial cells were isolated from breast cancer tissue specimens, and co-cultured with autologous MOs, isolated from peripheral blood mononuclear cells (PBMCs). First, tumor cells were cultured alone to check the MET effects on both proliferation and viability using BrdU (Bromodeoxyuridine [5-bromo-2’-deoxyuridine]), and Trypan Blue Exclusion Test [TBET], respectively, and on p-Akt-to-Akt ratios. Similarly, MOs were cultured alone for phagocytosis capacity assays. LDH-based cytotoxicity, respiratory burst and redox activity (nitric oxide [NO], catalase, superoxide dismutase [SOD]), release of antitumor cytokine IFN-γ, and immunosuppressive/regulatory cytokine IL-10, inducible nitric oxide synthase iNOS-associated proinflammatory MOs and arginase activities-associated anti-inflammatory MOs, and intracellular free calcium ions (_if_Ca^2+^) were measured in MOs cultured alone and co-cultured with breast cancer cells. All experiments were repeated four times (Figure 1).

**Figure 1.**
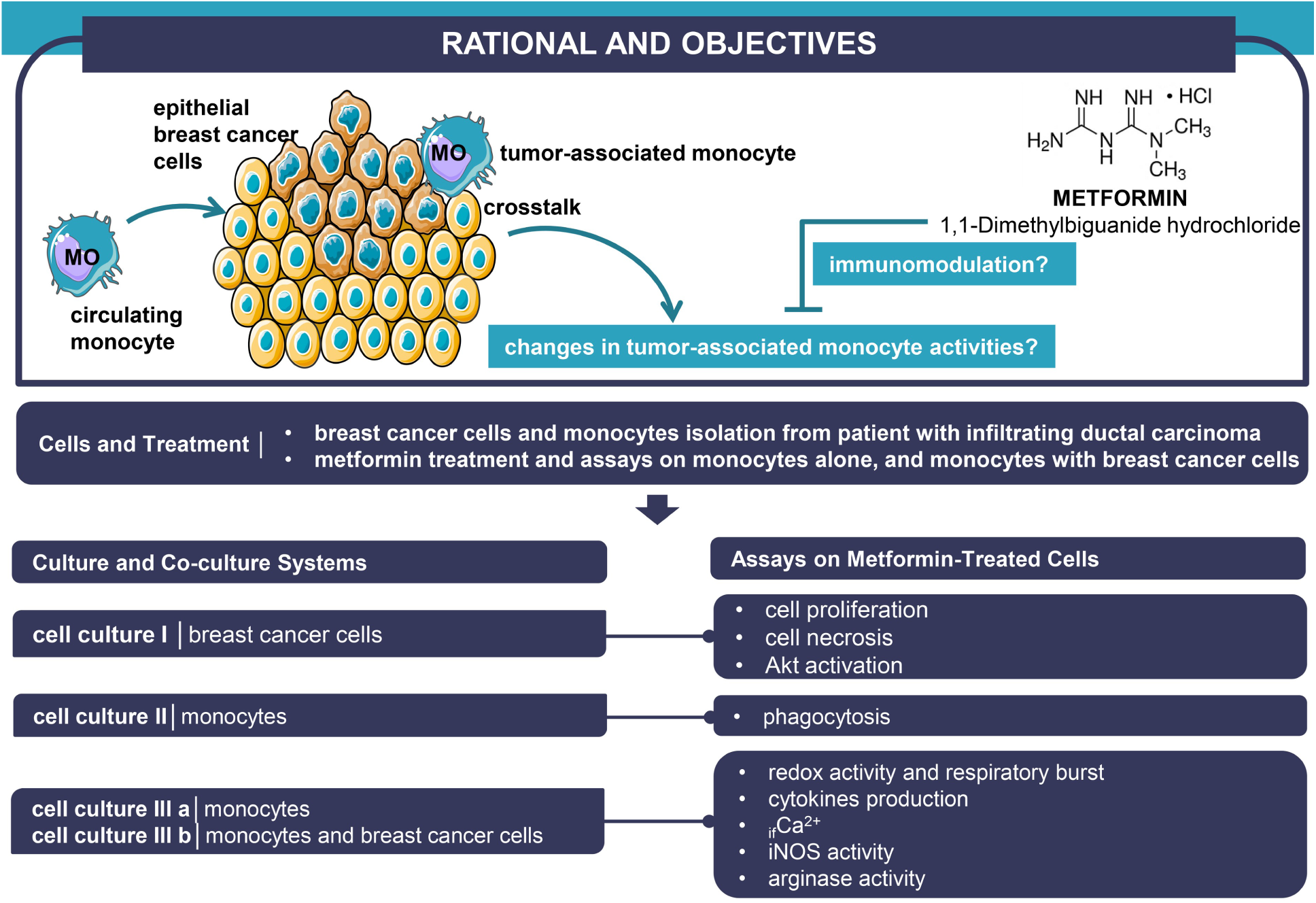
Summary of experimental design. Akt: protein kinase B (PKB), _if_Ca^2+^: intracellular free calcium ions, iNOS: nitric oxide synthase, MO: monocyte.

#### 2. Specimen samples

Primary tumor tissue specimens and autologous peripheral blood samples were collected thanks to three patient volunteers with breast cancer (age group 50-60 years), admitted to the Surgery Department of Tlemcen Medical Centre University (Algeria), after obtaining informed and written consent from each to participate to the current study. Peripheral blood was collected in heparinized *Vacutainer* tubes (BD, Belliver Industrial Estate, UK). The homogeneity of sample biopsies and absence of intertumor variations for each tumor intended for experimental analyses were checked by macroscopical and thorough anatomopathological examinations, and based on cellular morphology and immunohistochemical analysis. All tumor samples are characterized by the absence of estrogen receptor (ER) and the presence of human epidermal growth factor receptor 2 (HER2+). The current study was approval by local Ethics Committee of Tlemcen University, in accordance with the Declaration of Helsinki.

#### 3. Isolation of mammary adherent tumor epithelial cells

After removal of the healthy tissue surrounding the tumor tissue, tumor mammary epithelial cells presenting as infiltrating ductal carcinomas were isolated from primary cancer specimens by enzymatic digestion and differential centrifugation according to Feller et al., and Speirs et al., (17,18), with some modifications. Briefly, the tumor tissue specimens were washed extensively with 1x phosphate saline buffer (PBS), placed in sterile Petri dishes and cut into small 2 mm pieces with a sterile scalpel. The minced tissue was incubated in 0.1% collagenase solution at 37 °C for 12-20 h. Following digestion, cell mixtures were centrifuged at 40 x *g* for 1 min and the supernatant transferred to new tubes that were then centrifuged at 100 x *g* for 2 min to obtain a pellet representing tumor epithelial cells.

#### 4. Cell culture

Epithelial cancer cells were washed with RPMI-1640 medium, supplemented with 10% fetal calf serum (FCS) and 50 μg/mL gentamicin, by centrifugation at 40 x *g* for 5 min. The cell pellet was resuspended with 10 mL of RPMI-1640 supplemented with 10% FCS and 50 μg/mL gentamicin and then subdivided into two culture flasks and incubated in a humidified atmosphere at 37 °C and 5% CO_2_. Fibroblast contamination was removed from epithelial cancer cells by differential trypsinization (19).The culture medium was changed every 2-5 days (20). Cells were passaged with 0.25% trypsin-EDTA (ethylenediamine tetraacetic acid) when they reached ∼80% confluence (21).

#### 5. Peripheral blood mononuclear cells isolation

Blood samples were diluted 1:1 with PBS and layered on Histopaque-1077 (Sigma-Aldrich, St. Louis MO, USA) and centrifuged at 400 x *g* for 30 min. The interface band containing PBMCs was carefully harvested washed twice with PBS. Cell pellets were suspended in 1 mL of RPMI-1640 supplemented with 10% FCS and 50 μg/mL of gentamicin for cell counting. Cell viability was performed by TBET using photonic microscopy (Zeiss, Germany).

#### 6. MOs isolation

MOs were isolated from PBMCs based on differential plastic adherence (22). Briefly, PBMCs were cultivated in RPMI-1640 supplemented with 10% FCS and 50 μg/mL gentamicin, and seeded at 2 × 10^6^ cell/mL into 24-well plates. Cells were allowed to adhere for 2 h at 37 °C before removal of non-adherent cells were and treatment of adherent MOs with MET. Cells were counted microscopically (Zeiss, Germany) using trypan blue staining, and the purity of monocytes was evaluated by fluorescent staining with PhycoErytherin (PE)-anti-human CD14 antibody (BD Biosciences, San Diego, CA, USA) using a Floid Cell Imaging Station (Thermo Fisher Scientific, MA USA) (23,24) and routinely exceeded over 90% purity.

#### 7. Cell culture and co-culture systems

After cell detachment with trypsin-EDTA (25), breast cancer cells were counted before being cultured alone or co-cultured with MOs at a ratio of 1:1 in RPMI-1640 supplemented with 10% FCS and 50 μg/mL gentamicin.

#### 8. MET treatment

MOs, breast cancer cells or co-cultured MOs with breast cancer cells were treated for 24 h with fresh medium containing or not MET at the dose of 2.5 mM (15).

#### 9. TBET assays

The effect of MET treatment on cancer cell viability was based on TBET (trypan blue exclusion test). Breast cancer cells (2 × 10^5^ cells per well) were grown overnight in a 24-well plate at 37 °C in a humidified atmosphere and 5% CO_2_ for adherence. Thereafter, culture medium was replaced with fresh RPMI-1640 medium containing different concentrations of MET and incubated a further 24 h. Cells were subsequently washed with 1 x PBS, trypsinized before determination the number of viable and dead cells with TBET.

#### 10. BrdU assays

Cell proliferation was measured by BrdU incorporation using a BrdU Cell Proliferation ELISA according to the manufacturer’s instructions (ab126556-BrdU Cell Proliferation kit, Abcam, Germany). Briefly, breast cancer cells (2 × 10^5^ cells/mL) were treated with MET in a 96-well microplates for 24 h at 37 °C in a humidified atmosphere and 5% CO_2_. Thereafter, 20 µL of the diluted 1x BrdU was added to each well and cells were incubated overnight. Cells were then fixed and BrdU incorporation detected using anti-BrdU monoclonal Detector Antibody for 1 h at room temperature before incubation with peroxidase goat anti-mouse IgG conjugate as secondary antibody. Color was developed using tetramethylbenzidine (TMB) as a peroxidase substrate and BrdU incorporation measured at 450 nm using an ELISA reader (Biochrom Anthos 2020, Cambridge, UK).

#### 11. LDH-based cytotoxicity assays

Lactate dehydrogenase (LDH)-based cytotoxicity levels were determined by evaluation of LDH release into the cell culture supernatants using Lactate Dehydrogenase Activity Assay kit (MAK066, Sigma-Aldrich). Briefly, 50 μL of supernatant and 50 μL of the Master Reaction Mix were mixed and added to each well of a 96-well plate. Absorbance was measured at 450 mn after 30 min incubation at 37 °C in accordance with the manufacturer’s instructions.

#### 12. Cytokine assays

Concentration of anti-tumor-associated cytokine IFN-γ and immunosuppressive-associated cytokine IL-10 in cell culture media of MOs or co-culture system supernatants were measured by sandwich enzyme-linked immunosorbent assays (ELISA), using respective commercial kits (BD Biosciences), according to the manufacturer’s instructions. Optical densities (OD) were measured at 450 nm using appropriate standard curves for each cytokine.

#### 13. Arginase activity assays

The enzymatic activity of arginase (EC 3.5.3.1) was evaluated in cell lysates based on determination of urea levels following the L-arginine hydrolysis as described in detail (26). The arginase activity was expressed as mU urea/mg protein/1 h.

#### 14. _if_Ca^2+^ assay

The concentrations of _if_Ca^2+^ were measured biochemically based on the orthocresolphthalein complexone method as described elsewhere (27).

#### 15. Redox activity and respiratory burst assays

Redox activity and respiratory burst were evaluated by determination of the levels of NO production, catalase and SOD activities.

##### 15.1 NO assay and iNOS activity

The accumulation of NO in cell-free culture supernatants were evaluated by nitrite (NO_2_) measurement, as stable and final end-product of NO, using a sensitive colorimetric Griess reaction as described (28). Absorbance was measured at 540 nm using a Biochrom Anthos 2020 ELISA plate reader. NO production levels were calculated by comparison with a sodium nitrite (NaNO_2_) curve standard (29). iNOS activity was obtained by normalizing each NO concentration to milligrams of protein and expressed as picomoles/mg protein/30 min.

##### 15.2 Catalase activity assay

The enzymatic activity of catalase was spectrophotometrically determined in cell lysates by measurement of hydrogen peroxide (H_2_O_2_) decomposition (30). 10 μL volumes of cell lysates were added to a reaction mixture of H_2_O_2_ in 0.9% (v/v) aqueous saline before incubation for 5 min. The reactions were stopped by the addition of Titanyl sulfate (TiOSO_4_), and the absorbance measured at 410 nm.

##### 15.3 SOD activity assay

SOD (superoxide dismutase) activity in cell lysates was determined spectrophotometrically by measuring production of a water-soluble formazan dye resulting from the reduction of Dojindo’s highly water-soluble tetrazolium salt WST-1 ((2-4-Iodophenyl)-3-(4-nitrophenyl)-5-(2,4-disulfophenyl)-2H-tetrazolium, monosodium salt), using a SOD Assay Kit-WST (19160, Sigma Aldrich). Twenty μL of the enzyme working solution were added to a mixture containing 20 μL of cell lysate and 200 μL of WST Working Solution. The microplate was incubated at 37 °C for 20 min, and the absorbance read at 440 nm. The SOD activity (percentage inhibition of WST-1 inhibition) was calculated as follows:

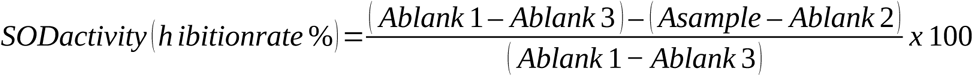

#### 16. Phagocytosis assay

Assay for phagocytosis capacity was performed as described (30–32). Briefly, a total of 2 × 10^5^ MOs were infected with *Staphylococcus aureus* at multiplicity of infection (MOI) of 50 in a 24-well plate and incubated with different concentrations of MET for 1 h at 37 °C in a 5% CO_2_ incubator. The number of viable bacteria was determined by serial dilution and colony forming unit (CFU) counts on Chapman medium. The percentage of phagocytosis was calculated as follows:

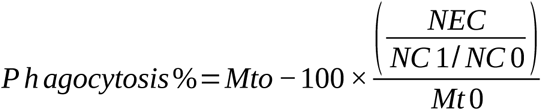

M_t0_ is the number of bacteria in the assay sample mixture at t_0_. NEC, number of extracellular bacteria in assay mixtures sample at t_1_. NC_0_ and NC_1_ correspond to control sample at t_0_ and t_1_.

#### 17. Western blotting assays

After 24 h incubation of breast cancer cells treated or not with MET, cells were washed with PBS and lysed using Triton X-100. Proteins present in equal amounts of cell lysates were rapidly diluted with SDS-sample buffer (50 mM Tris-HCL pH 6.8, 2 mM DTT, 1.0% SDS), boiled for 5 min. Proteins were separated by 10% sodium dodecyl-sulfate polyacrylamide gel electrophoresis (SDS-PAGE). Protein concentrations were not determined before reduction and denaturation to minimize the chance of protein dephosphorylation. After separation, proteins were transferred to a nitrocellulose membranes and transferred protein were visualized by staining with Ponceau red. Thereafter, membranes were blocked with 5% nonfat milk or 5% bovine serum albumin (BSA) for 45 min at room temperature and incubated overnight at 4 °C with primary antibodies against p-Akt (Ser473) (1/1000), Akt-1 (2H10) (1/1000), Akt-2 (5B5) (1/1000) Cell Signaling Technology (Denvers, MA, USA). Horseradish peroxidase-conjugated (HRP) anti-mouse IgG and anti-rabbit IgG were used as secondary antibodies for 1 h at room temperature. Blotted membranes were detected with enhanced chemiluminescence reagent (Amersham Pico) using X-ray film. Quantitative analysis of the signals from scanned films was performed using Imgcalc2, a unix software developed in house (IGH, Montpellier) for quantifying pixels on numerical images. The results in Figure 2c-d are represented as a ratio to the signal in Ponceau Red staining to correct for differences in total protein loading with the levels for MET at dose 0 set as 1.

**Figure 2.**
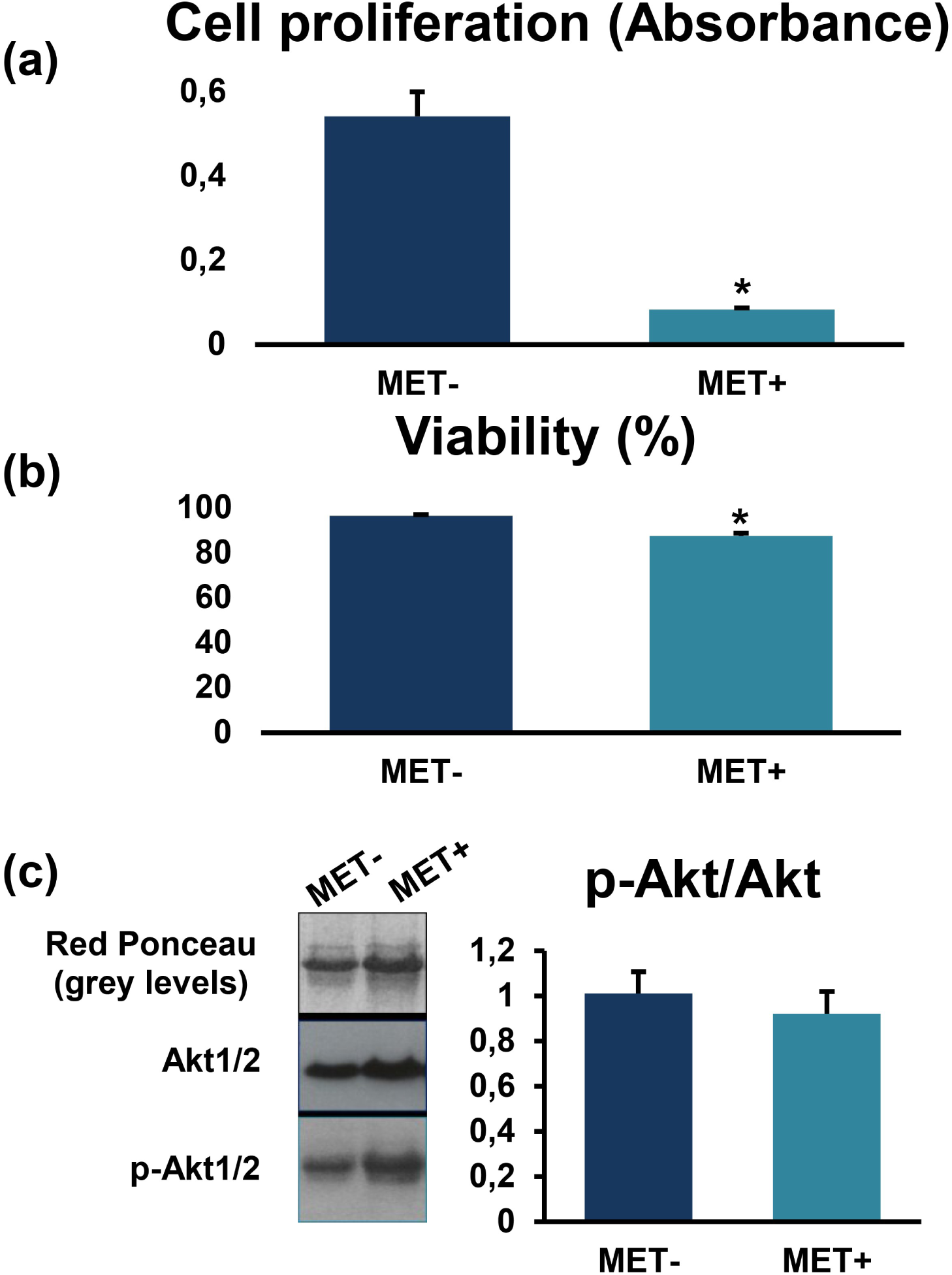
Effect of MET treatment on breast cancer cell proliferation, viability and the ratio of phosphorylated Akt-to-Akt. (a) In breast cancer cells treated or not with MET, cell proliferation was determined by BrdU assay and (b) viability by TBET assay. (c) Levels of major proteins stained by red Ponceau are shown to use as loading control. Values are represented as a ratio to the protein levels in Ponceau red and with the value for zero MET set as 1. MET: metformin, p-Akt: phosphorylated Akt. Asterisks indicate significant differences when comparing treated group with untreated controls (0 mM MET) using Mann-Whitney *U* test (\**p* < 0.05).

#### 18. Statistical analysis

Data are presented as the mean with standard errors of means (SEM). Statistical analyses were performed using non-parametric Mann-Whitney *U* or Kruskal-Wallis one-way analysis of variance (ANOVA) test with pairwise comparisons using the Dunn–Bonferroni approach after checking the distribution of data. Statistics were carried out using IBM SPSS Statistics version 20. *P*-values less than 0.05 were considered significant.

## Results

### MET downregulates breast cancer cell viability and proliferation, and the ratio of phosphorylated Akt1/2 *versus* total Akt1/2

As shown in Figures 2 (a) and (b), MET treatment significantly downregulated breast cancer cell proliferation and viability levels (for both comparisons, *p* < 0.05). Figure 2 (c) shows the raw data of the actual protein levels (upper panels), total Akt1/2 levels (middle panels) and levels of phospho-(activated) Akt1/2 (lower panels). The histogram shows the relative expression levels of activated Akt1/2-to-total Akt1/2 ratio after normalization to the total protein loaded. As observed in Figure 2 (c), MET treatment did not show a significant difference in the Akt levels when comparing with MET-untreated cells, but the ratio of activated Akt *versus* total Akt was downregulated; nevertheless, the difference did not reach the significance level (*p* > 0.05). So the actions of MET on Akt1/2 are essentially due to a loss/reduction of Akt1/2 activity since when the ratio of active Akt1/2 (phosphorylated) to total Akt1/2 levels is calculated we observed a reduction of Akt1/2 activity in cancer cells.

### MET induces upregulation of necrosis in co-culture system of breast cancer cells and MOs, but has no cytotoxic effect on MOs cultured alone

As indicated in Figure 3, MET treatment induced no necrosis/LDH-based cytotoxicity effects on MO cells (*p* > 0.05). Additionally, the levels of LDH-based cytotoxicity were significantly upregulated in MET-treated co-culture systems. Conversely, the level of LDH-based cytotoxicity was significantly downregulated in MET-untreated co-cultures of MOs with breast cancer cells when compared to MET-untreated MOs cultured alone (*p* < 0.05).

**Figure 3.**
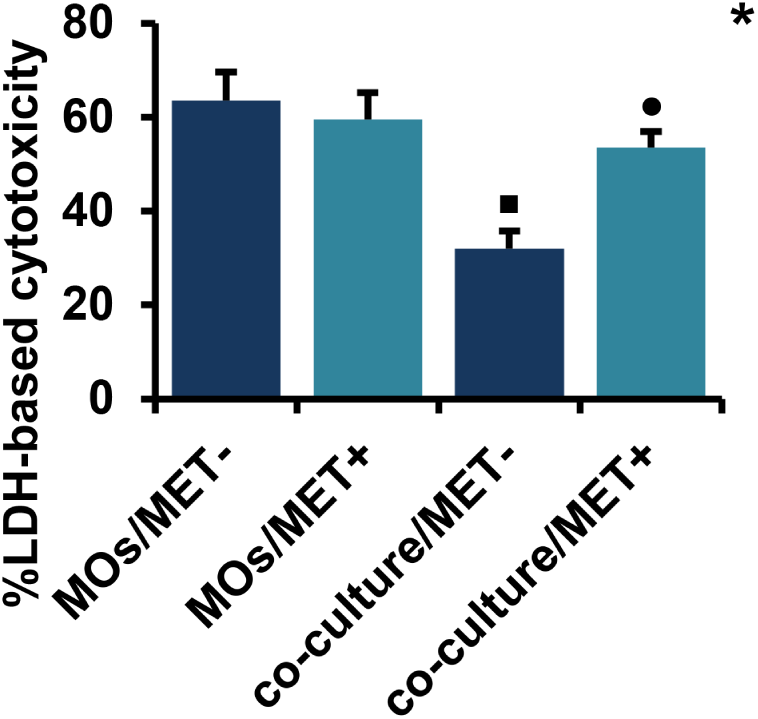
Cytotoxic effect of MET on MOs and co-cultured cells. Necrosis levels were measured spectrophotometrically through the evaluation of LDH release. Values are presented as the mean with standard error of mean for four independent experiments carried out on three samples (n = 12 for each group). MET: metformin, LDH: lactate dehydrogenase, MOs: monocytes, MOs/MET-: MET-untreated MOs, MOs/MET+: MET-treated MOs, co-culture/MET-: MET-untreated co-culture system, co-culture/MET+: MET-treated co-culture system. Black dots indicate significant differences when comparing each treated group with untreated controls (0 mM MET) using Mann-Whitney *U* test (•*p* < 0.05). Black boxes indicate significant differences highlighted between MET-untreated MOs and MET-untreated co-culture system using Mann-Whitney *U* test (□*p* < 0.05). Asterisks indicate significant differences highlighted between all groups by Kruskal-Wallis test with pairwise Dunn-Bonferroni adjustment (\**p* < 0.05).

### iNOS and arginase activities are simultaneously upregulated in co-cultures of MOs with breast cancer cells after treatment with MET

As depicted in Figure 4, MET induced an increase in iNOS activity in co-culture systems, as compared to MET-treated or MET-untreated MOs cultured alone; while the difference was not significant for the comparison with MET-treated MOs (respectively, *p* > 0.05 and *p* < 0.05). Additionally, iNOS activity was significantly upregulated in MET-untreated MOs co-cultured with breast cancer cells in comparison to MET-untreated MOs cultured alone (*p* < 0.05). Similarly, MET significantly increased the arginase activity of MOs cultured alone or co-cultured with breast cancer cells (*p* < 0.05). However, arginase activity was significantly downregulated in MET-untreated MOs co-cultured with breast cancer cells compared to MET-untreated MOs cultured alone (*p* < 0.05).

**Figure 4.**
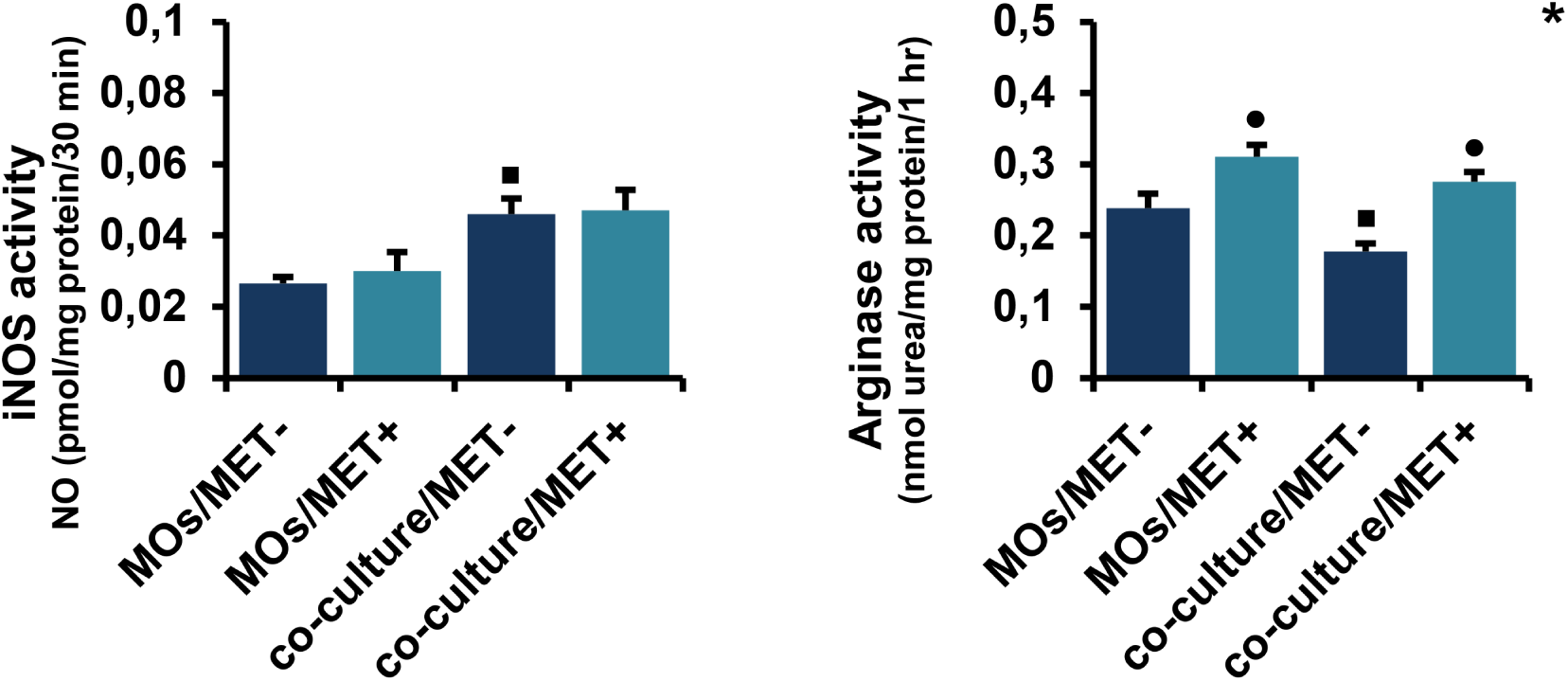
Effect of MET on iNOS and arginase activities in MOs and co-culture system. NO levels were measured by Griess colorimetric reaction and iNOS activity was obtained by normalizing each NO to protein concentrations and time. The enzymatic activity of arginase was evaluated in cell lysates by the spectrophotometric measurement of urea concentration. Values are presented as the mean with standard error of mean for four independent experiments carried out on three samples (n = 12 for each group). MET: metformin, MOs: monocytes, NO: nitric oxide, iNOS: inducible nitric oxide synthase, MOs/MET-: MET-untreated MOs, MOs/MET+: MET-treated MOs, co-culture/MET-: MET-untreated co-culture system, co-culture/MET+: MET-treated co-culture system. Black dots indicate significant differences when comparing each treated group with untreated controls (0 mM MET) using Mann-Whitney *U* test (•*p* < 0.05). Black boxes indicate significant differences highlighted between MET-untreated MOs and MET-untreated co-culture system using Mann-Whitney *U* test (□*p* < 0.05). Asterisks indicate significant differences highlighted between all groups by Kruskal-Wallis test with pairwise Dunn-Bonferroni adjustment (\**p* < 0.05).

### MET downregulated Monocyte phagocytosis

As shown in Figure 5, the phagocytic activity of MOs significantly decreased after MET treatment (*p* < 0.05).

**Figure 5.**
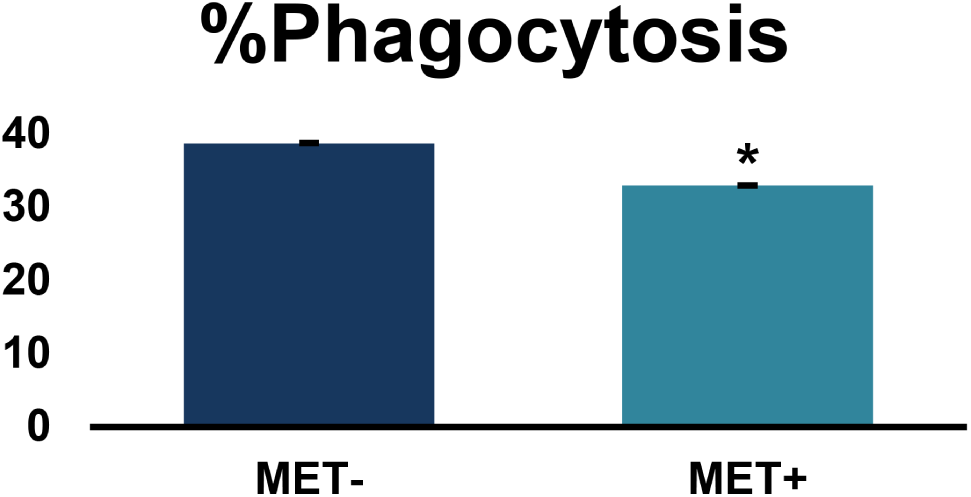
Effect of MET on MOs phagocytosis capacity. MOs were infected with *Staphylococcus aureus* before treatment with MET. The results were expressed as a percentage of phagocytosis. Values are presented as the mean with standard error of mean for four independent experiments carried out on three samples (n = 12 for each group). MET: metformin, MOs: monocytes. Asterisks indicate significant differences between treated cells and untreated controls by Mann-Whitney *U* test (\**p* < 0.05).

### MET improves redox activity

As demonstrated in Figure 6, MET had no significant effect on catalase activity in MO cells while inducing a significant increase of catalase activity in co-culture systems (*p* < 0.05). Additionally, catalase activity was significantly downregulated in MET-untreated co-cultures of MOs with breast cancer cells than in MET-untreated MOs cultured alone (*p* < 0.05). Moreover, SOD activity was significantly increased in MET-treated compared to MET-untreated MOs (*p* < 0.05). In contrast to MOs cultured alone, MET had no significant effect on SOD activity in co-culture systems.

**Figure 6.**
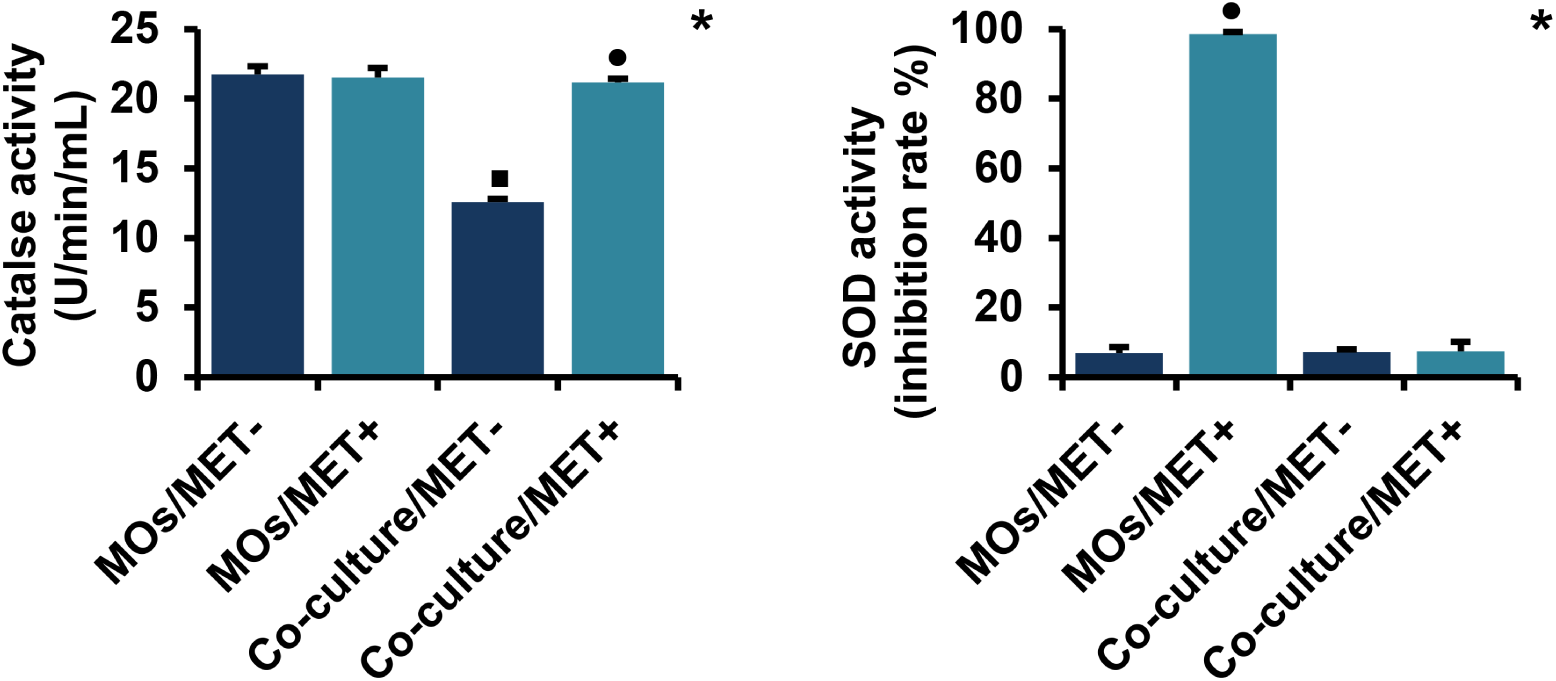
Effect of MET on catalase and SOD activities in MOs and co-culture system. Catalase activity was determined spectrophotometrically by measurement of hydrogen peroxide decomposition. SOD activity was evaluated by spectrophotometric measurement of a water-soluble formazan dye. Values are presented as the mean with standard error of mean for four independent experiments carried out on three samples (n = 12 for each group). MET: metformin, MOs: monocytes, SOD: superoxide dismutase, MOs/MET-: MET-untreated MOs, MOs/MET+: MET-treated MOs, co-culture/MET-: MET-untreated co-culture system, co-culture/MET+: MET-treated co-culture system. Black dots indicate significant differences when comparing each treated group with untreated controls (0 mM MET) using Mann-Whitney *U* test (•*p* < 0.05). Black boxes indicate significant differences highlighted between MET-untreated MOs and MET-untreated co-culture system using Mann-Whitney *U* test (□*p* < 0.05). Asterisks indicate significant differences highlighted between all groups by Kruskal-Wallis test with pairwise Dunn-Bonferroni adjustment (\**p* < 0.05).

### The effects of MET on _if_Ca^2+^ differs between MOs cultured alone and MOs co-cultured with breast cancer cells

As shown in Figure 7, MET had no significant effect on _if_Ca^2+^ levels in MOs cells while inducing a significant increase of _if_Ca^2+^ levels in MOs co-cultured with breast cancer cells (*p* < 0.05). Conversely, MET treatment downregulated _if_Ca^2+^ in MET-untreated MOs co-cultivated with breast cancer cells more significantly than in MET-untreated MOs cultured alone (*p* < 0.05).

**Figure 7.**
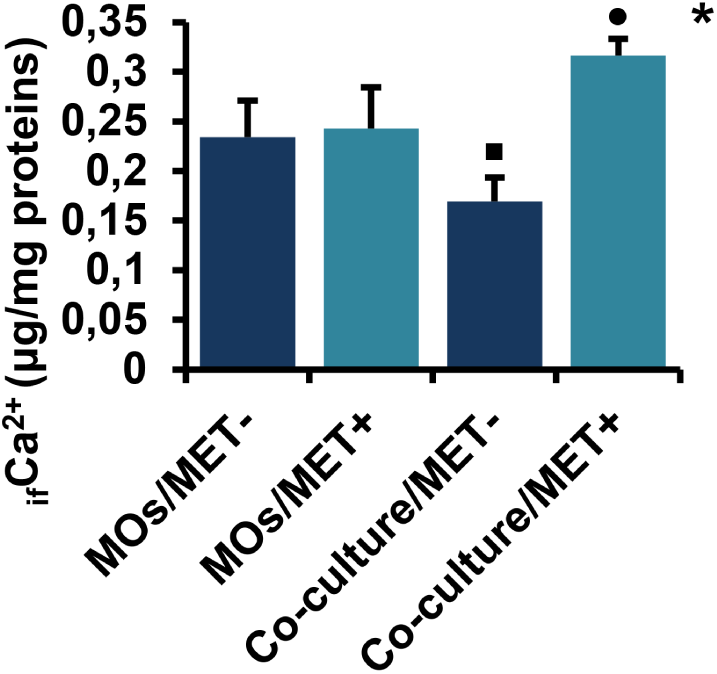
Effect of MET on _if_Ca^2+^ levels in MOs and co-culture system. Values are presented as the mean with standard error of mean for four independent experiments carried out on three samples (n = 12 for each group). MET: metformin, MOs: monocytes, _if_Ca^2+^: intracellular free calcium ions, MOs/MET-: MET-untreated MOs, MOs/MET+: MET-treated MOs, co-culture/MET-: MET-untreated co-culture system, co-culture/MET+: MET-treated co-culture system. Black dots indicate significant differences when comparing each treated group with untreated controls (0 mM MET) using Mann-Whitney *U* test (•*p* < 0.05). Black boxes indicate significant differences highlighted between MET-untreated MOs and MET-untreated co-culture system using Mann-Whitney *U* test (□*p* < 0.05). Asterisks indicate significant differences highlighted between all groups by Kruskal-Wallis test with pairwise Dunn-Bonferroni adjustment (\**p* < 0.05).

### MET enhances production of IFN-γ in both MOs and co-culture systems, but downregulates the production of IL-10 in MOs cultured alone

As shown in Figure 8, MET induced a significant upregulation of IFN-γ levels in both MOs and MOs co-cultured with breast cancer cells in co-culture systems (for both comparisons, *p* < 0.05). Conversely, MET treatment induced significant downregulation of IL-10 production in MOs while inducing upregulation in co-cultured cells, but the difference was not significant (respectively, *p* < 0.05 and *p* > 0.05). Moreover, IFN-γ and IL-10 levels were decreased significantly in MET-untreated MOs co-cultured with breast cancer cells when compared to MET-untreated MOs cultured alone (for both comparisons, *p* < 0.05).

**Figure 8.**
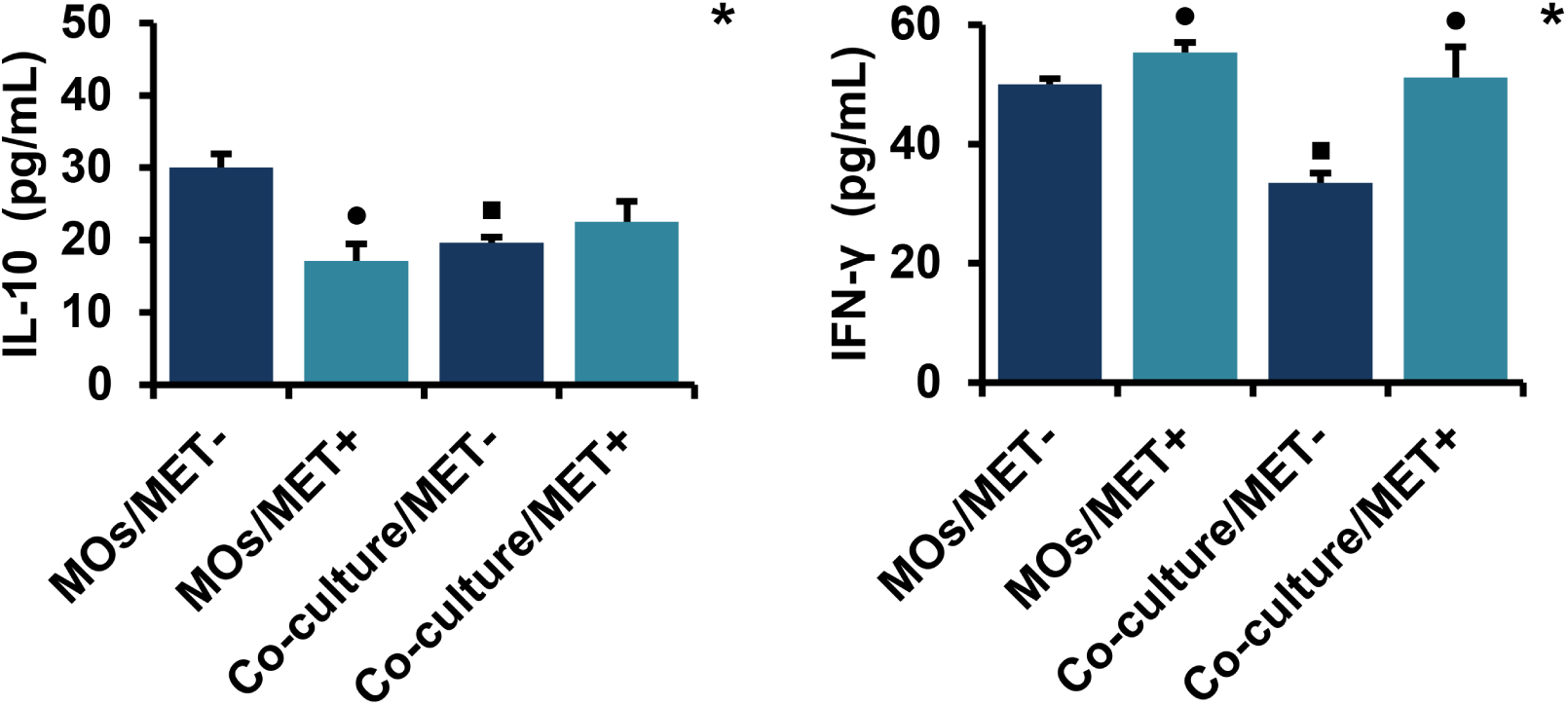
Effect of MET on the production of IL-10 and IFN-γ in MOs and co-culture system. IL-10 and IFN-γ levels were measured using sandwich enzyme-linked immunosorbent assay (ELISA). Values are presented as the mean with standard error of mean for four independent experiments carried out on three samples (n = 12 for each group). MET: metformin, MOs: monocytes, IFN: interferon, IL: interleukin, MOs/MET-: MET-untreated MOs, MOs/MET+: MET-treated MOs, co-culture/MET-: MET-untreated co-culture system, co-culture/MET+: MET-treated co-culture system. Black dots indicate significant differences when comparing each treated group with untreated controls (0 mM MET) using Mann-Whitney *U* test (•*p* < 0.05). Black boxes indicate significant differences highlighted between MET-untreated MOs and MET-untreated co-culture system using Mann-Whitney *U* test (□*p* < 0.05). Asterisks indicate significant differences highlighted between all groups by Kruskal-Wallis test with pairwise Dunn-Bonferroni adjustment (\**p* < 0.05).

## Discussion

MET has recently received increasing attention as a potential therapeutic treatment against cancer (34). Here we have examined the effects of MET in a novel co-culture system comprising primary MOs and breast cancer cells. Measuring several different parameters including Akt activation, cell necrosis, iNOS and arginase activities, cellular redox, calcium signaling and cytokine release, we confirm scientific relevance of the co-culture system over the use of isolated cell types for analyzing the differential effects of MET in a tumor-like microenvironment.

The dose of MET used in these experiments (2.5 mM) is relatively higher than those used clinically for the treatment of type 2 diabetes (ca 0.5 mM). However, it is important to note that *in vitro* cultured cells are maintained under less physiological conditions. In particular, cultured cells do not benefit from the indirect anti-tumor effects of MET occurring *in vivo* such as the reduction of insulin levels - where insulin is known to have a mitogenic effect - and cultured cells are exposed to high concentrations of growth factors and glucose present in the culture medium, which may help explain the required higher doses of MET (35).

The anti-tumorigenic properties of MET have been reported in several studies associated with indirect action (reduced insulin levels) or direct actions on molecular pathways that regulate breast tumor cell growth and death (36). MET may mediate its effects through actions on different cells of the tumor microenvironment, including MOs-macrophages that would be involved in controlling tumor cell growth and progression. However, the activities of immune cells could undoubtedly change when they are in contact with tumor cells. In this context, we investigated the effect of MET on functional activities of autologous MOs cultured alone and when co-cultured with primary breast cancer cells. In conclusion, our results demonstrate a significant effect of MET on co-cultures of MOs with breast cancer cells on several indicators of anti-tumoral activity as summarized in Figure 9.

Here, we first tested the effects of MET on proliferation and viability of cancer cells. We found that MET downregulated both the proliferation and viability of breast cancer cells. Our results are consistent with those obtained recently using BrdU and 3-(4,5-Dimethylthiazol-2-yl)-2,5-diphenyltetrazolium bromidefor (MTT) assays on breast cancer cell lines MCF-7, MDA-MB-231 and MDA-MB-435 (15,37). The same effects were observed on MCF-7 and MDA-MB-231 cell viability with the doses of at 1 mM and 5 mM of MET using TBET assays after 24 h and 48 h of treatment (38). In terms of cell activation and proliferation, the PI3K/Akt/mTOR signaling pathway has been highlighted to play an important role in vital cell functions including cell growth, proliferation, differentiation and survival (39). Its hyper-activation can lead to excessive tumor cell proliferation, inhibition of apoptosis, angiogenesis, invasion and metastasis (40–42). For our part, we suggest that MET treatment may play a noticeable role in modulating cell signaling activation related to PI3K/Akt/mTOR pathways in breast cancer cells, that corroborates recent reports (43,44).

It is well known that necrosis and iNOS are both involved in tissue damage occurring during inflammation. Our results showed that MET treatment had no effect on MOs necrosis. However, a marked cytotoxic effect of MET was observed in co-culture systems, which is in agreement with previous *in vitro* studies, carried out on BT-20 breast cancer cell line (45). In MOs, as well as in other cells especially macrophages, the amino acid L-arginine is also used as a substrate by arginase to produce polyamines (46) that contribute to the tumor progression (47), and by iNOS to produce NO (46), which has antitumor effects at high levels (48). The current study provides evidence that MOs cultured with breast cancer cells exhibited high levels of iNOS, but remain without marked change when treated with MET. Our observations are in agreement with earlier findings demonstrating that MET induces antitumoral activity of macrophages during breast cancer (49) and suppresses polarization toward pro-tumoral phenotypes (50). Although arginase activity was downregulated in untreated co-cultured cells, it was upregulated after MET treatment. Hence, MET has been reported to have opposing effects in normal or pathological conditions whereby it can both attenuate NO production and enhance arginase activation in MOs and macrophages (51,52). It would be of interest to check the impact of MET treatment on iNOS and arginase activities simultaneously.

The phagocytic activity of MOs is the subject of several studies under normal and pathological conditions including breast cancer (53). In our study, we observed that pretreatment with MET downregulated the phagocytic capacity of MOs, which is of interest, knowing that high phagocytic capacity is associated with MOs that promote survival and extravasation of cancer cells, and characterizes the so-called ‘classical MOs’ in humans (54,55).

The link between cancer and altered metabolism has previously been suggested as a common feature of cancerous tissues, such as the Warburg effect, in which some antioxydant molecules, like NADPH, can be used in protective mechanisms against oxidative stress and ROS that are produced during rapid cell proliferation (56). High levels of ROS can cause macromolecular damage, that can lead to apoptosis and senescence (57). Our findings demonstrated that catalase activity was downregulated in MET-untreated co-cultures, but no differences were observed when comparing MET-treated co-cultures and both MET-untreated or MET-treated MOs cultured alone. Additionally, SOD activity was changed only in MET-treated MOs. In summary, MET treatment did not show metabolic alterations with regard to the levels of the antioxidant molecules catalase and SOD within the co-cultures of MOs with breast cancer cells compared to MOs cultivated alone.

_if_Ca^2+^ is an important secondary messenger that regulates various cellular processes and signaling pathways including those related to cancer, such as apoptosis, proliferation and metastasis (58,59), and those involved in immune responses of MOs, including the production of cytokines and phagocytic activation (60,61). We first observed that _if_Ca^2+^ levels were reduced in MOs when co-cultured with breast cancer cells. In constrast, MET treatment induced _if_Ca^2+^ upregulation in both MOs and co-culture systems. Our findings support simultaneous MO activation and the induction of apoptosis in breast cancer cells as suggested previously (59,62,63).

It is now accepted that IL-10 enhances cancer immune surveillance and suppression of cancer-associated inflammation (64), as well as inducing expression of IFN-γ (65) which exerts anti-tumor activities directly by enhancement of tumor cells antigenicity, inhibition of cell proliferation, the induction of apoptosis or indirectly by inhibition of angiogenesis (66). Our results indicate that the levels of both IL-10 and IFN-γ decreased during interplay between MOs and breast cancer cells. Treatment with MET has shown differences between its action when MOs are cultured alone and when co-cultured with breast cancer cells. MET induced a decrease in IL-10 levels in MOs and, conversely, an increase in IFN-γ levels. However, MET treatment induced upregulation in the production of both cytokines when the MOs were co-cultured with breast cancer cells, although the differences were not significant for IL-10. Our results demonstrate that Metformin treatment is clearly effective and important in a system that shares some similarities with the biological system where mononuclear cells are not alone but may be confronted with malignant cells.

## Conclusions and Future Prospects

*In fine*, our results not only show that the activities of human MOs change when they interact with autologous primary breast cancer cells, but also provide the first evidence that MET treatment upregulates the production of antitumor IFN-γ and downregulates the proliferation and viability of breast cancer cells, as well as improves cytotoxicity during MOs-breast cancer cells crosstalk. These findings open the route to further investigations including the study of MET on autophagy/reverse Warburg effects, as well as the molecular characterization of different subpopulations of monocytes involved in the interaction with breast cancer cells following MET treatment.

## Acknowledgments

The authors are grateful to the Team W0414101 of the Laboratory of Applied Molecular Biology and Immunology (University of Tlemcen, Algeria) for their help and technical assistance. They would also like to express their deep gratitude to the Thematic Research Agencies ATRBSA and ATRSS (Algeria) and the Directorate General of Scientific Research and Technological Development (DGRSDT, Direction Générale de la Recherche Scientifique et du Développement Technologique, MESRS, Algeria) for their financial support.

## References

1. Ghoncheh M, Pournamdar Z, Salehiniya H. Incidence and Mortality and Epidemiology of Breast Cancer in the World. Asian Pac J Cancer Prev APJCP (2016) 17:43–46.

2. Soliman H. Immunotherapy strategies in the treatment of breast cancer. Cancer Control J Moffitt Cancer Cent (2013) 20:17–21.

3. Gingras I, Azim HA, Ignatiadis M, Sotiriou C. Immunology and breast cancer: toward a new way of understanding breast cancer and developing novel therapeutic strategies. Clin Adv Hematol Oncol HO (2015) 13:372–382.

4. Sabatier R, Finetti P, Mamessier E, Adelaide J, Chaffanet M, Ali HR, Viens P, Caldas C, Birnbaum D, Bertucci F. Prognostic and predictive value of PDL1 expression in breast cancer. Oncotarget (2015) 6:5449–5464. doi:10.18632/oncotarget.3216

5. Xu T, He B-S, Liu X-X, Hu X-X, Lin K, Pan Y-Q, Sun H-L, Peng H-X, Chen X-X, Wang S-K. The Predictive and Prognostic Role of Stromal Tumor-infiltrating Lymphocytes in HER2-positive Breast Cancer with Trastuzumab-based Treatment: a Meta-analysis and Systematic Review. J Cancer (2017) 8:3838–3848. doi:10.7150/jca.21051

6. Doseff A, Parihar A. “Monocyte Subsets and Their Role in Tumor Progression,” in Tumor Microenvironment and Myelomonocytic Cells, ed. S. Biswas (InTech). Available at: http://www.intechopen.com/books/tumor-microenvironment-and-myelomonocytic-cells/monocyte-subsets-and-their-role-in-tumor-progression [Accessed September 27, 2017]

7. Ben-Baruch A. Host microenvironment in breast cancer development: Inflammatory cells, cytokines and chemokines in breast cancer progression: reciprocal tumor– microenvironment interactions. Breast Cancer Res (2002) 5: doi:10.1186/bcr554

8. Evani SJ, Prabhu RG, Gnanaruban V, Finol EA, Ramasubramanian AK. Monocytes mediate metastatic breast tumor cell adhesion to endothelium under flow. FASEB J (2013) 27:3017–3029. doi:10.1096/fj.12-224824

9. Mohamed M, Cavallo-Medved D, Rudy D, Anbalagan A, Moin K, Sloane B. Interleukin-6 Increases Expression and Secretion of Cathepsin B by Breast Tumor-Associated Monocytes. Cell Physiol Biochem (2010) 25:315–324. doi:10.1159/000276564

10. Chiang C-F, Chao T-T, Su Y-F, Hsu C-C, Chien C-Y, Chiu K-C, Shiah S-G, Lee C-H, Liu S-Y, Shieh Y-S. Metformin-treated cancer cells modulate macrophage polarization through AMPK-NF-&#x3BA;B signaling. Oncotarget (2017) 8: doi:10.18632/oncotarget.14982

11. Stout RD, Suttles J. Functional plasticity of macrophages: reversible adaptation to changing microenvironments. J Leukoc Biol (2004) 76:509–513. doi:10.1189/jlb.0504272

12. Fan C, Wang Y, Liu Z, Sun Y, Wang X, Wei G, Wei J. Metformin exerts anticancer effects through the inhibition of the Sonic hedgehog signaling pathway in breast cancer. Int J Mol Med (2015) 36:204–214. doi:10.3892/ijmm.2015.2217

13. Liu B, Fan Z, Edgerton SM, Yang X, Lind SE, Thor AD. Potent anti-proliferative effects of metformin on trastuzumab-resistant breast cancer cells via inhibition of erbB2/IGF-1 receptor interactions. Cell Cycle (2011) 10:2959–2966. doi:10.4161/cc.10.17.16359

14. Camacho L, Dasgupta A, Jiralerspong S. Metformin in breast cancer - an evolving mystery. Breast Cancer Res (2015) 17: doi:10.1186/s13058-015-0598-8

15. Queiroz EAIF, Puukila S, Eichler R, Sampaio SC, Forsyth HL, Lees SJ, Barbosa AM, Dekker RFH, Fortes ZB, Khaper N. Metformin Induces Apoptosis and Cell Cycle Arrest Mediated by Oxidative Stress, AMPK and FOXO3a in MCF-7 Breast Cancer Cells. PLoS ONE (2014) 9:e98207. doi:10.1371/journal.pone.0098207

16. Ding L, Liang G, Yao Z, Zhang J, Liu R, Chen H, Zhou Y, Wu H, Yang B, He Q. Metformin prevents cancer metastasis by inhibiting M2-like polarization of tumor associated macrophages. Oncotarget (2015) 6: doi:10.18632/oncotarget.5541

17. Feller WF, Stewart SE, Kantor J. Primary tissue culture explants of human breast cancer. J Natl Cancer Inst (1972) 48:1117–1120.

18. Speirs V, Green AR, Walton DS, Kerin MJ, Fox JN, Carleton PJ, Desai SB, Atkin SL. Short-term primary culture of epithelial cells derived from human breast tumours. Br J Cancer (1998) 78:1421–1429.

19. Jones JCR. Reduction of contamination of epithelial cultures by fibroblasts. CSH Protoc (2008) 2008:pdb.prot4478. doi:10.1101/pdb.prot4478

20. Farnie G, Clarke RB, Spence K, Pinnock N, Brennan K, Anderson NG, Bundred NJ. Novel Cell Culture Technique for Primary Ductal Carcinoma In Situ: Role of Notch and Epidermal Growth Factor Receptor Signaling Pathways. JNCI J Natl Cancer Inst (2007) 99:616–627. doi:10.1093/jnci/djk133

21. Benton G, DeGray G, Kleinman HK, George J, Arnaoutova I. In Vitro Microtumors Provide a Physiologically Predictive Tool for Breast Cancer Therapeutic Screening. PLOS ONE (2015) 10:e0123312. doi:10.1371/journal.pone.0123312

22. Wahl LM, Wahl SM, Smythies LE, Smith PD. “Isolation of Human Monocyte Populations,” in Current Protocols in Immunology, eds. J. E. Coligan, B. E. Bierer, D. H. Margulies, E. M. Shevach, W. Strober (Hoboken, NJ, USA: John Wiley & Sons, Inc.). Available at: http://doi.wiley.com/10.1002/0471142735.im0706as70 [Accessed September 30, 2017]

23. Pabst MJ, Pabst KM, Handsman DB, Beranova-Giorgianni S, Giorgianni F. Proteome of monocyte priming by lipopolysaccharide, including changes in interleukin-1beta and leukocyte elastase inhibitor. Proteome Sci (2008) 6:13. doi:10.1186/1477-5956-6-13

24. Zhou L, Somasundaram R, Nederhof RF, Dijkstra G, Faber KN, Peppelenbosch MP, Fuhler GM. Impact of human granulocyte and monocyte isolation procedures on functional studies. Clin Vaccine Immunol CVI (2012) 19:1065–1074. doi:10.1128/CVI.05715-11

25. Sarvaiya HA, Lazar IM. Insulin stimulated MCF7 breast cancer cells: Proteome dataset. Data Brief (2016) 9:579–584. doi:10.1016/j.dib.2016.09.025

26. Aribi M. Macrophage Bactericidal Assays. Methods Mol Biol Clifton NJ (2018) 1784:135–149. doi:10.1007/978-1-4939-7837-3_14

27. Takano S, Kaji H, Hayashi F, Higashiguchi K, Joukei S, Kido Y, Takahashi J, Osawa K. A calculation model for serum ionized calcium based on an equilibrium equation for complexation. Anal Chem Insights (2012) 7:23–30. doi:10.4137/ACI.S9681

28. Kavoosi G, Ardestani SK, Kariminia A, Tavakoli Z. Production of nitric oxide by murine macrophages induced by lipophosphoglycan of Leishmania major. Korean J Parasitol (2006) 44:35. doi:10.3347/kjp.2006.44.1.35

29. Blond D, Raoul H, Le Grand R, Dormont D. Nitric Oxide Synthesis Enhances Human Immunodeficiency Virus Replication in Primary Human Macrophages. J Virol (2000) 74:8904–8912. doi:10.1128/JVI.74.19.8904-8912.2000

30. Walton PA. Effects of peroxisomal catalase inhibition on mitochondrial function. Front Physiol (2012) 3: doi:10.3389/fphys.2012.00108

31. Aribi M, Meziane W, Habi S, Boulatika Y, Marchandin H, Aymeric J-L. Macrophage Bactericidal Activities against Staphylococcus aureus Are Enhanced In Vivo by Selenium Supplementation in a Dose-Dependent Manner. PLOS ONE (2015) 10:e0135515. doi:10.1371/journal.pone.0135515

32. Herrera MT, Gonzalez Y, Hernández-Sánchez F, Fabián-San Miguel G, Torres M. Low serum vitamin D levels in type 2 diabetes patients are associated with decreased mycobacterial activity. BMC Infect Dis (2017) 17: doi:10.1186/s12879-017-2705-1

33. Nouari W, Ysmail-Dahlouk L, Aribi M. Vitamin D3 enhances bactericidal activity of macrophage against Pseudomonas aeruginosa. Int Immunopharmacol (2016) 30:94–101. doi:10.1016/j.intimp.2015.11.033

34. Zhuang Y, Miskimins WK. Metformin Induces Both Caspase-Dependent and Poly(ADP-ribose) Polymerase-Dependent Cell Death in Breast Cancer Cells. Mol Cancer Res (2011) 9:603–615. doi:10.1158/1541-7786.MCR-10-0343

35. Garofalo C, Capristo M, Manara MC, Mancarella C, Landuzzi L, Belfiore A, Lollini P-L, Picci P, Scotlandi K. Metformin as an adjuvant drug against pediatric sarcomas: hypoxia limits therapeutic effects of the drug. PloS One (2013) 8:e83832. doi:10.1371/journal.pone.0083832

36. Xin W, Fang L, Fang Q, Zheng X, Huang P. Effects of metformin on survival outcomes of pancreatic cancer patients with diabetes: A meta-analysis. Mol Clin Oncol (2018) 8:483–488. doi:10.3892/mco.2017.1541

37. Gao Z-Y, Liu Z, Bi M-H, Zhang J-J, Han Z-Q, Han X, Wang H-Y, Sun G-P, Liu H. Metformin induces apoptosis via a mitochondria-mediated pathway in human breast cancer cells in vitro. Exp Ther Med (2016) 11:1700–1706. doi:10.3892/etm.2016.3143

38. Marinello PC, da Silva TNX, Panis C, Neves AF, Machado KL, Borges FH, Guarnier FA, Bernardes SS, de-Freitas-Junior JCM, Morgado-Díaz JA, et al. Mechanism of metformin action in MCF-7 and MDA-MB-231 human breast cancer cells involves oxidative stress generation, DNA damage, and transforming growth factor β1 induction. Tumor Biol (2016) 37:5337–5346. doi:10.1007/s13277-015-4395-x

39. Mendoza MC, Er EE, Blenis J. The Ras-ERK and PI3K-mTOR pathways: cross-talk and compensation. Trends Biochem Sci (2011) 36:320–328. doi:10.1016/j.tibs.2011.03.006

40. Miller TW, Hennessy BT, González-Angulo AM, Fox EM, Mills GB, Chen H, Higham C, García-Echeverría C, Shyr Y, Arteaga CL. Hyperactivation of phosphatidylinositol-3 kinase promotes escape from hormone dependence in estrogen receptor–positive human breast cancer. J Clin Invest (2010) 120:2406–2413. doi:10.1172/JCI41680

41. Mutlu M, Saatci Ö, Ansari SA, Yurdusev E, Shehwana H, Konu Ö, Raza U, Sahin Ö. miR-564 acts as a dual inhibitor of PI3K and MAPK signaling networks and inhibits proliferation and invasion in breast cancer. Sci Rep (2016) 6: doi:10.1038/srep32541

42. Sadeghi N, Gerber DE. Targeting the PI3K pathway for cancer therapy. Future Med Chem (2012) 4:1153–1169. doi:10.4155/fmc.12.56

43. Alimova IN, Liu B, Fan Z, Edgerton SM, Dillon T, Lind SE, Thor AD. Metformin inhibits breast cancer cell growth, colony formation and induces cell cycle arrest in vitro. Cell Cycle (2009) 8:909–915. doi:10.4161/cc.8.6.7933

44. Al-Zaidan L, El Ruz RA, Malki AM. Screening Novel Molecular Targets of Metformin in Breast Cancer by Proteomic Approach. Front Public Health (2017) 5:277. doi:10.3389/fpubh.2017.00277

45. Szewczyk M, Richter C, Briese V, Richter D-U. A retrospective in vitro study of the impact of anti-diabetics and cardioselective pharmaceuticals on breast cancer. Anticancer Res (2012) 32:2133–2138.

46. Rath M, MÃ¼ller I, Kropf P, Closs EI, Munder M. Metabolism via Arginase or Nitric Oxide Synthase: Two Competing Arginine Pathways in Macrophages. Front Immunol (2014) 5: doi:10.3389/fimmu.2014.00532

47. Avtandilyan N, Javrushyan H, Petrosyan G, Trchounian A. The Involvement of Arginase and Nitric Oxide Synthase in Breast Cancer Development: Arginase and NO Synthase as Therapeutic Targets in Cancer. BioMed Res Int (2018) 2018:1–9. doi:10.1155/2018/8696923

48. Keshet R, Erez A. Arginine and the metabolic regulation of nitric oxide synthesis in cancer. Dis Model Mech (2018) 11:dmm033332. doi:10.1242/dmm.033332

49. Chiang C-F, Chao T-T, Su Y-F, Hsu C-C, Chien C-Y, Chiu K-C, Shiah S-G, Lee C-H, Liu S-Y, Shieh Y-S. Metformin-treated cancer cells modulate macrophage polarization through AMPK-NF-kB signaling. Oncotarget (2017) 8: doi:10.18632/oncotarget.14982

50. Ding L, Liang G, Yao Z, Zhang J, Liu R, Chen H, Zhou Y, Wu H, Yang B, He Q. Metformin prevents cancer metastasis by inhibiting M2-like polarization of tumor associated macrophages. Oncotarget (2015) 6: doi:10.18632/oncotarget.5541

51. Buldak L, Labuzek K, Buldak RJ, Kozlowski M, Machnik G, Liber S, Suchy D, Dulawa-Buldak A, Okopień B. Metformin affects macrophages’ phenotype and improves the activity of glutathione peroxidase, superoxide dismutase, catalase and decreases malondialdehyde concentration in a partially AMPK-independent manner in LPS-stimulated human monocytes/macrophages. Pharmacol Rep (2014) 66:418–429. doi:10.1016/j.pharep.2013.11.008

52. Kato Y, Koide N, Komatsu T, Tumurkhuu G, Dagvadorj J, Kato K, Yokochi T. Metformin Attenuates Production of Nitric Oxide in Response to Lipopolysaccharide by Inhibiting MyD88-Independent Pathway. Horm Metab Res (2010) 42:632–636. doi:10.1055/s-0030-1255033

53. Baskić D, Aćimović LD, Arsenijević N. [The phagocytic activity of monocytes in different stages of breast cancer]. Med Pregl (2003) 56 Suppl 1:103–107.

54. Cassetta L, Pollard JW. Cancer immunosurveillance: role of patrolling monocytes. Cell Res (2016) 26:3–4. doi:10.1038/cr.2015.144

55. Al Dubayee MS, Alayed H, Almansour R, Alqaoud N, Alnamlah R, Obeid D, Alshahrani A, Zahra MM, Nasr A, Al-Bawab A, et al. Differential Expression of Human Peripheral Mononuclear Cells Phenotype Markers in Type 2 Diabetic Patients and Type Diabetic Patients on Metformin. Front Endocrinol (2018) 9:537. doi:10.3389/fendo.2018.00537

56. Zahzeh MR, Loukidi B, Meziane W, Haddouche M, Mesli N, Zouaoui Z, Aribi M. Relationship between NADPH and Th1/Th2 ratio in patients with non-Hodgkin lymphoma who have been exposed to pesticides. J Blood Med (2015) 6:99–107. doi:10.2147/JBM.S78759

57. Cairns RA, Harris IS, Mak TW. Regulation of cancer cell metabolism. Nat Rev Cancer (2011) 11:85–95. doi:10.1038/nrc2981

58. Monteith GR, Davis FM, Roberts-Thomson SJ. Calcium Channels and Pumps in Cancer: Changes and Consequences. J Biol Chem (2012) 287:31666–31673. doi:10.1074/jbc.R112.343061

59. Muhammad SNH, Mokhtar NF, Yaacob NS. 15d-PGJ2 Induces Apoptosis of MCF-7 and MDA-MB-231 Cells via Increased Intracellular Calcium and Activation of Caspases, Independent of ERα and ERβ. Asian Pac J Cancer Prev APJCP (2016) 17:3223–3228.

60. Brown D, Donaldson K, Stone V. Effects of PM10 in human peripheral blood monocytes and J774 macrophages. Respir Res (2004) 5: doi:10.1186/1465-9921-5-29

61. Gronski MA, Kinchen JM, Juncadella IJ, Franc NC, Ravichandran KS. An essential role for calcium flux in phagocytes for apoptotic cell engulfment and the anti-inflammatory response. Cell Death Differ (2009) 16:1323–1331. doi:10.1038/cdd.2009.55

62. Cross BM, Breitwieser GE, Reinhardt TA, Rao R. Cellular calcium dynamics in lactation and breast cancer: from physiology to pathology. Am J Physiol-Cell Physiol (2014) 306:C515–C526. doi:10.1152/ajpcell.00330.2013

63. França E, Honorio-França A, Nunes G, Fagundes D, Marchi P, Fernandes R, França J, Botelho A, Varotti F, Moraes L. Intracellular calcium is a target of modulation of apoptosis in MCF-7 cells in the presence of IgA adsorbed to polyethylene glycol. OncoTargets Ther (2016)617. doi:10.2147/OTT.S99839

64. Dennis KL, Blatner NR, Gounari F, Khazaie K. Current status of interleukin-10 and regulatory T-cells in cancer. Curr Opin Oncol (2013) 25:637–645. doi:10.1097/CCO.0000000000000006

65. Mumm JB, Emmerich J, Zhang X, Chan I, Wu L, Mauze S, Blaisdell S, Basham B, Dai J, Grein J, et al. IL-10 elicits IFNγ-dependent tumor immune surveillance. Cancer Cell (2011) 20:781–796. doi:10.1016/j.ccr.2011.11.003

66. Mojic M, Takeda K, Hayakawa Y. The Dark Side of IFN-γ: Its Role in Promoting Cancer Immunoevasion. Int J Mol Sci (2017) 19:89. doi:10.3390/ijms19010089

